# Pre-exposure to mRNA-LNP inhibits adaptive immune responses and alters innate immune fitness in an inheritable fashion

**DOI:** 10.1101/2022.03.16.484616

**Authors:** Zhen Qin, Aurélie Bouteau, Christopher Herbst, Botond Z. Igyártó

## Abstract

Hundreds of millions of SARS-CoV-2 mRNA-LNP vaccine doses have already been administered to humans. However, we lack a comprehensive understanding of the immune effects of this platform. The mRNA-LNP-based SARS-CoV-2 vaccine is highly inflammatory, and its synthetic ionizable lipid component responsible for the induction of inflammation has a long *in vivo* half-life. Since chronic inflammation can lead to immune exhaustion and non-responsiveness, we sought to determine the effects of pre-exposure to the mRNA-LNP on adaptive immune responses and innate immune fitness. We found that pre-exposure to mRNA-LNPs or LNP alone led to long-term inhibition of the adaptive immune responses, which could be overcome using standard adjuvants. On the other hand, we report that after pre-exposure to mRNA-LNPs, the resistance of mice to heterologous infections with influenza virus increased while *Candida albicans* decreased. The diminished resistance to *Candida albicans* correlated with a general decrease in blood neutrophil percentages. Interestingly, mice pre-exposed to the mRNA-LNP platform can pass down the acquired immune traits to their offspring, providing better protection against influenza. In summary, the mRNA-LNP vaccine platform induces long-term unexpected immunological changes affecting both adaptive immune responses and heterologous protection against infections. Thus, our studies highlight the need for more research to determine this platform’s true impact on human health.

**Authors Summary:** We bring experimental evidence that pre-exposure to mRNA-LNPs or its LNP component affects innate and adaptive immune responses. Pre-exposure to mRNA-LNPs led to long-term inhibition of the adaptive immune responses, which the use of adjuvants could overcome. On the other hand, we report that after pre-exposure to mRNA-LNPs, the resistance of mice to heterologous infections with influenza virus increased while *Candida albicans* decreased. We also detected a general neutropenia in the mRNA-LNP exposed mice. Interestingly, mice pre-exposed to mRNA-LNPs can pass down the acquired immune traits to their offspring. In summary, the mRNA-LNP vaccine platform induces long-term immunological changes that can affect both adaptive immune responses and heterologous protection against infections, some of which can be inherited by the offspring. More studies are needed to understand the mechanisms responsible for these effects and determine this platform’s impact on human health.

## INTRODUCTION

The mRNA-LNP vaccine platform gained much attention with the ongoing SARS-CoV-2 pandemic. Initially, this vaccine platform was thought to be non-inflammatory since the mRNA has been modified and purified to limit innate immune activation [1–3]. At the same time, the lipid nanoparticle (LNP) component was considered an inert carrier and protector of the mRNA. However, recently has been shown that the synthetic ionizable lipid component of the LNPs is highly inflammatory [4], and this is a critical component to support induction of adaptive immune responses. These LNPs mixed with proteins induce comparable responses to mRNA-LNPs [5]. The platform can support the induction of adaptive immune responses in the absence of a variety of different inflammatory cytokines, -pathways, and innate immune cells [5–7].

The acute side effects reported with the mRNA-LNP vaccine platform are diverse and likely associated with its highly inflammatory nature and partially mediated by innate immune responses [4,8]. In addition to the induction of specific T- and B-cell activation, certain vaccines or infections can affect long-term innate immune responses by either increasing or decreasing the activation of innate immune cells [9]. Furthermore, the innate immune reprogramming induced by certain vaccines can interfere with the immune responses induced by other vaccines [9]. The possible short and long-term immunological changes of the mRNA-LNP vaccine outside the induction of antigen-specific anti-SARS-CoV-2 responses are unknown. A recent human study awaiting peer-review reported innate and adaptive immune reprogramming with this platform [10], while single-cell RNA-seq studies on human white blood cells derived from vaccinated people also revealed significant changes in innate immune cells [11]. Whether the reported changes are long-lasting and can influence immune fitness or interfere with the responses induced by other vaccines remains to be determined.

Here, using an mRNA-LNP animal vaccination model, we show that pre-exposure to mRNA-LNP inhibits antibody responses. The inhibition could be overcome with the use of adjuvants and did not interfere with protein vaccines efficacy. At the same time however, this vaccine platform enhances innate immune fitness towards influenza infection but decreases resistance to *Candida albicans*. The enhanced immune fitness towards influenza can be passed down to the offspring.

## RESULTS

### Pre-exposure to LNPs or mRNA-LNPs inhibit adaptive immune responses

The LNPs used in preclinical animal studies are highly inflammatory [4]. The critical inflammatory component of the LNPs is the synthetic ionizable lipid, which for the Pfizer SARS-CoV-2 vaccine has been estimated to have a 20–30-day *in vivo* half-life [12]. The LNPs used for preclinical studies and the Pfizer vaccine are similar and produced by Acuitas Therapeutics [4–6]. The immune system under chronic stimulation often responds with exhaustion and non-responsiveness [13]. Since the mRNA-LNP platform is highly inflammatory and has a long *in vivo* half-life, we sought to test whether pre-exposure to this platform affects subsequent adaptive immune responses. We used an intradermal immunization model developed in our laboratory [4,6] to test this. Adult WT mice were exposed to PBS, 2.5 μg of mRNA-LNPs coding for eGFP, or 2.5 μg empty LNPs intradermally, as shown in **Figure 1A**. Two weeks later, the mice were injected in the same area with 2.5 μg of mRNA-LNP coding for PR8 influenza hemagglutinin (HA). Two weeks post-inoculation, the anti-HA responses were determined in the serum using ELISA and the GC B cell responses in the skin draining lymph nodes monitored by flow cytometry (Supplemental Figure 1), as we previously described [6]. We found that pre-exposure to mRNA coding for an irrelevant protein (eGFP) or empty LNPs significantly decreased the anti-HA responses, both on antibody and GC B cell levels (**Figure 1B and C**). We found no difference between mRNA-LNP and empty LNP groups (**Figure 1B and C**). Thus, these data suggest that pre-exposure to this platform can inhibit subsequent adaptive immune responses and that the LNPs play a critical role in this.

**Figure 1.**
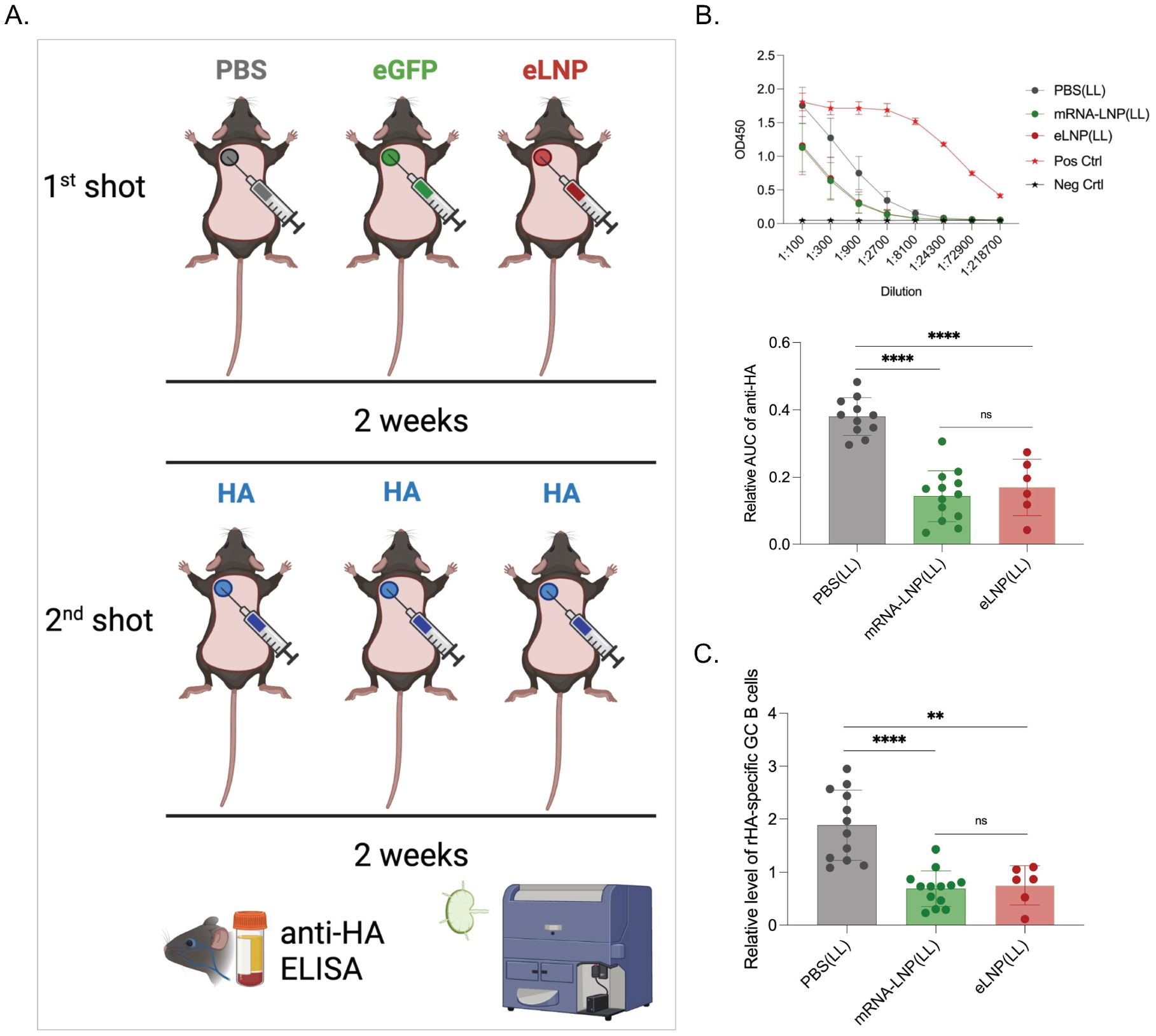
Pre-exposure to mRNA-LNPs or LNPs significantly inhibits subsequent adaptive immune responses induced by the mRNA-LNP vaccine. **A**). Experimental model. Animals were shaved and intradermally inoculated in the left upper spot with either PBS, mRNA-LNP coding for eGFP (eGFP) or empty LNP (eLNP). Two weeks later the same areas were injected with PR8 HA mRNA-LNP (HA). Serum and skin draining lymph nodes were harvested 2 weeks later and the anti-HA serum antibody levels determined using ELISA and the germinal center (GC) B cell responses using flow cytometer (please see Materials and Methods for details on data normalization). **B**). Serum anti-HA antibody levels detected by ELISA. OD450 readings (top) at different serum dilutions. Summary graph of the relative area under the curve (AUC) for each sample (middle). **C**). GC B cell responses (CD38^-^ GL7^+^) from the same mice. Each dot represents a separate mouse. Data from at least two separate experiments pooled and are shown as mean ±SD. One-way ANOVA was used to establish significance. ns = not significant. **p<0.005, ***p<0.0005, ****p<0.0001.

### The inhibition of adaptive immune responses by mRNA-LNPs is systemic, but more pronounced at the site of injection

Humans often receive follow-up and booster shots in the same arm with similar locations (deltoid muscle). In mice, the adaptive immune responses with the mRNA-LNP platform are primarily mounted in the lymph node draining the injection site [6]. Thus, we next sought to determine whether the inhibition we observed is localized or systemic. To define this, we performed similar experiments as detailed above, but one group of mice received the second shot in an area distinct from the first inoculation site (**Figure 2A**). We found that the mice injected in a different location still showed a significant decrease in anti-HA responses compared to PBS pretreatment (**Figure 2B and C**). However, the inhibition was significantly less pronounced than in mice injected into the same area (**Figure 2B and C**). Thus, pre-exposure to LNPs shows injection-site dominance with systemic traits.

**Figure 2.**
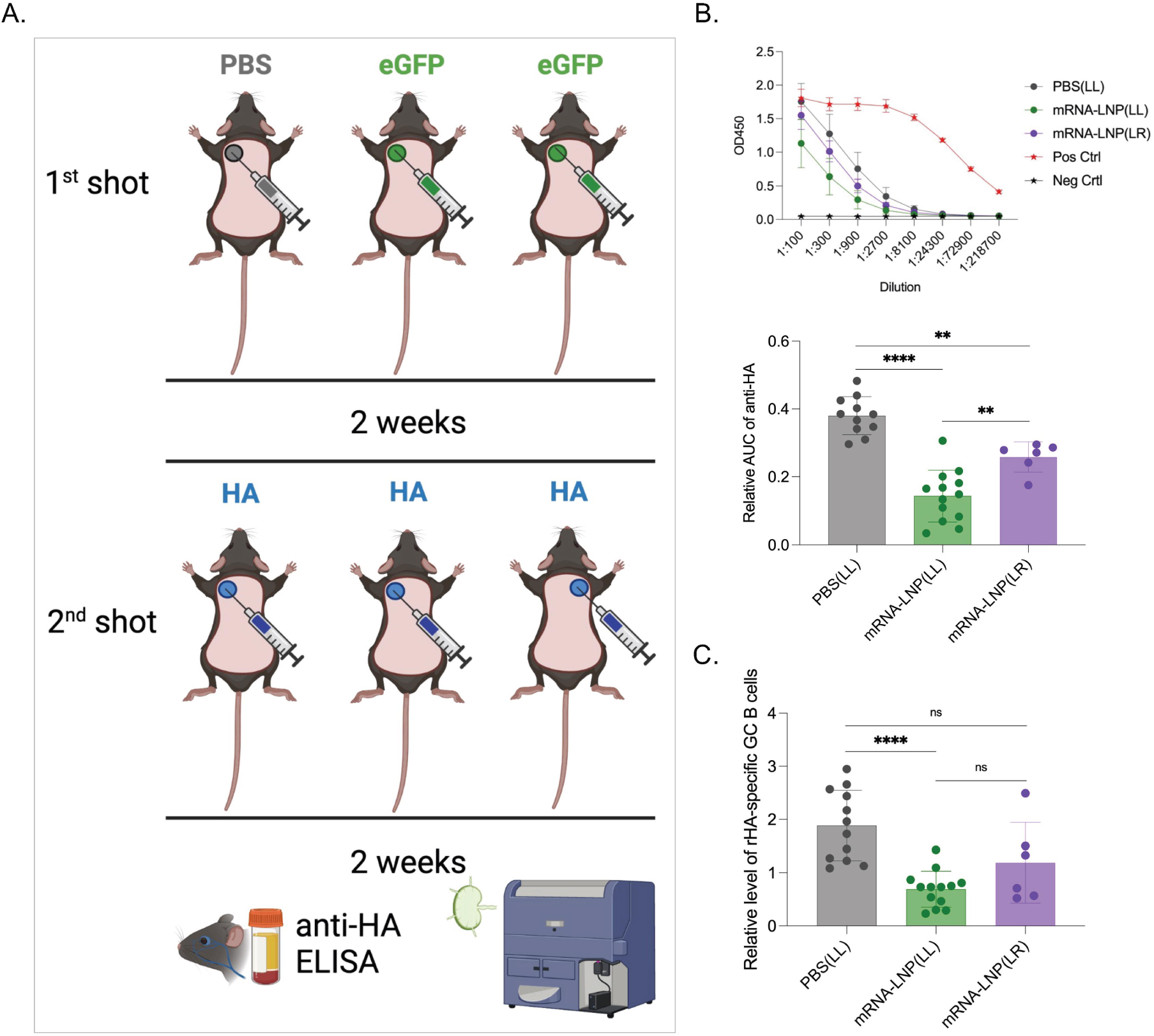
Pre-exposure to mRNA-LNPs has systemic effects. **A**). Experimental model. As in Figure 1, but one of the groups was injected at a spot distinct from the primary exposure. **B**). ELISA OD450 readings and summary AUC graph on serum anti-HA antibody levels. **C**). Relative GC B cell responses from the same mice. Each dot represents a separate mouse. Data from at least two separate experiments pooled and are shown as mean ±SD. One-way ANOVA was used to establish significance. ns = not significant. *p<0.05, **p<0.01, ****p<0.0001.

### Duration of the inhibitory effects of mRNA-LNPs on the adaptive immune responses

From a human health perspective, defining the length of the inhibitory effects caused by exposure to LNPs is critical to minimize its impact and devise preventive measures. To determine how long the inhibition lasted, WT mice were exposed to either PBS or mRNA-LNPs, and at 2, 4, or 8 weeks later injected into the same area with mRNA-LNPs coding for influenza HA as presented above (**Figure 3A**). Two weeks post-inoculation, we found that even mice injected four weeks post pre-exposure showed a significant decrease of anti-HA responses (**Figure 3B and C**). At week eight post-exposure, the GC B cell responses were still significantly lower in the mRNA-LNP exposed mice, but not the anti-HA antibody levels (**Figure 3B and C**). Thus, inhibition of the adaptive immune responses by pre-exposure to mRNA-LNPs is long-lasting but it is likely to wane with time.

**Figure 3.**
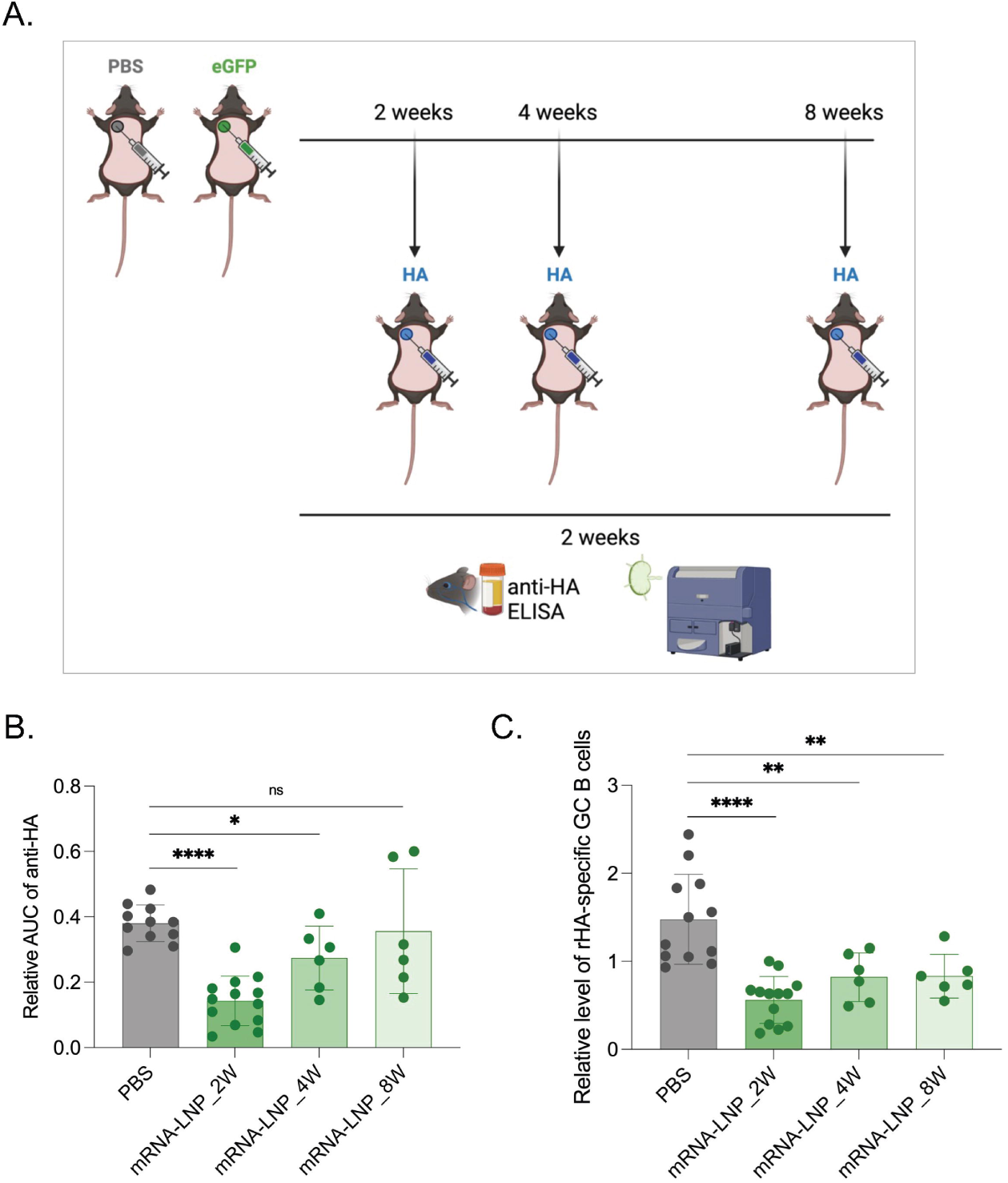
Pre-exposure to mRNA-LNPs has long-lasting effect on adaptive immune responses. **A**). Experimental model. Animals at the indicated timepoints post-exposure to PBS or eGFP mRNA-LNP were injected into the same spot with PR8 HA mRNA-LNP and then anti-HA responses assessed as depicted two weeks later. **B**). Summary AUC graph on serum anti-HA antibody levels. **C**). Relative GC B cell responses from the same mice. Each dot represents a separate mouse. Data from at least two separate experiments pooled and are shown as mean ±SD. One-way ANOVA was used to establish significance. ns = not significant. *p<0.05, **p<0.01, ****p<0.0001.

### Adjuvants might overcome the suppressive state induced by the pre-exposure to the mRNA-LNP platform

We observed that pre-exposure to LNPs significantly inhibited the adaptive immune responses triggered by the mRNA-LNP platform. However, it is crucial to determine whether there is a general inhibition of the adaptive immune responses, and whether pre-exposure to mRNA-LNP affects adaptive immune responses triggered by other vaccines, possibly altering immune protection. We first tested whether there is a generalized inhibition of adaptive immune responses with the pre-exposure to mRNA-LNP. For this, we used our well-established steady-state antigen targeting system that allows delivery of antigen to Langerin-expressing dendritic cells (DCs) with anti-Langerin antibody [14,15]. The antigen delivery to the DCs using this system does not induce activation and inflammation and has been successfully used to characterize the adaptive immune responses induced by DCs in steady-state [14,15]. As a first step, we adoptively transferred CFSE-labeled, congenically marked transgenic CD4 T cells specific to Eα peptide presented on IA_β_ into naïve WT B6 mice. The next day the mice were intradermally exposed to PBS or mRNA-LNP coding for GFP. On day 14 post-exposure, the mice were injected with PBS or 1 μg of anti-Langerin-Eα intravenously. Four days later, the mice were sacrificed, and the expansion of the TEα cells in the draining and non-draining LNs assessed using flow cytometry (**Figure 4A**). We found that the number of TEα cells in mice pre-exposed to mRNA-LNP, but received no Eα antigen, was lower than in PBS-treated mice (**Figure 4B**). In line with this, we observed that the expansion of the TEα cells was markedly reduced in both draining and non-draining LNs of the mRNA-LNP pre-exposed mice compared to the PBS-treated mice (**Figure 4B;** data from draining and non-draining LNs pooled). Thus, pre-exposure to mRNA-LNP inhibits T cell responses induced in a steady-state model, likely by decreasing precursor numbers. To determine whether the inhibition was on the T cell’s level, we repeated the experiments above, but the TEα cells were transferred 13 days after the mice were exposed to PBS or mRNA-LNP. In this setting, the transferred T cells, unlike the other host cells, such as DCs, are not exposed to mRNA-LNP, and the precursor numbers are expected to be similar between the groups. One day after transfer, the mice were inoculated with anti-Langerin-Eα, and the TEα cell expansion was determined 4 days later. In this setting, we detected normal expansion of TEα cells (**Figure 4B**). Thus, these data altogether suggest that the inhibition was on the T cell precursor levels.

**Figure 4.**
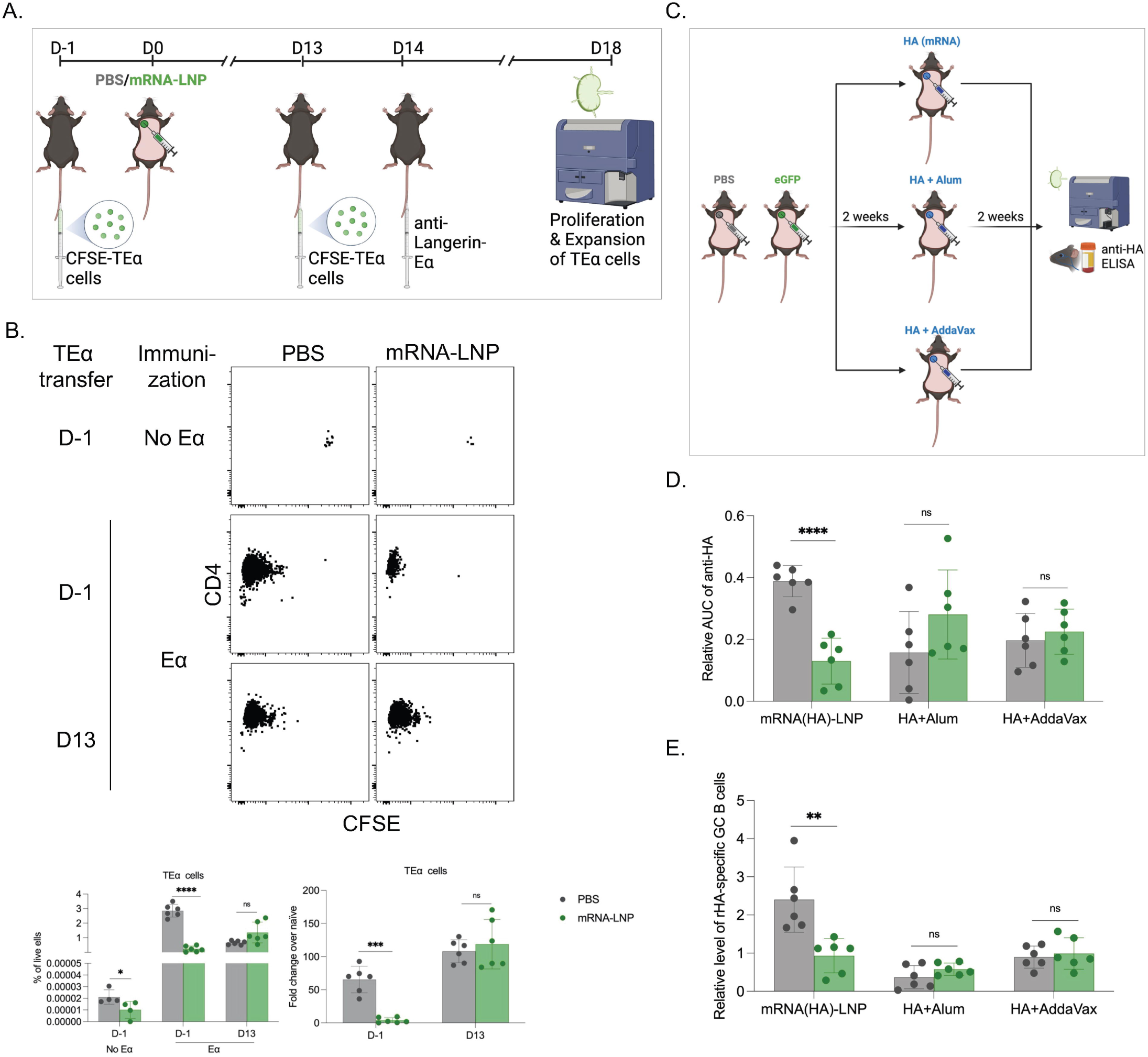
The general inhibition of adaptive immune responses induced by pre-exposure to mRNA-LNP can be overcome with the use of adjuvants. **A)**. Experimental model. CFSE-labeled TEα cells were transferred 1 day before (D-1) or 13 days after (D13) exposure to PBS or mRNA-LNP coding for eGFP. Anti-Langerin-Eα peptide was intravenously delivered on D14. The mice were sacrificed and the SDLNs were harvested for TEα cells examination by flow cytometry. **B**). Representative flow plots of TEα cell responses with/out Eα peptide stimulation (top), and the correspond summary graphs (bottom) on % of TEα cells in live cells and the fold change of expanded TEα cells over the non-immunized controls. **C**). Experimental model. Two weeks post-exposure to PBS or eGFP mRNA-LNP the animals were injected into the same spot with PR8 HA mRNA-LNP, or HA protein mixed with Alum or AddaVax. The anti-HA responses were assessed as depicted two weeks later. **D**). Summary AUC graph on serum anti-HA antibody levels. **E**). Relative GC B cell responses from the same mice. Each dot represents a separate mouse. Data from at least two separate experiments pooled and are shown as mean ±SD. Welch’s t test was used to establish significance. ns = not significant. **p<0.005, ***p<0.0005, ****p<0.0001.

To determine whether adjuvants can overcome the inhibition, PBS, and eGFP-coding mRNA-LNP pre-exposed mice were injected either with mRNA-LNPs coding for influenza PR8 HA or PR8 HA protein mixed with Alum (Th2 adjuvant) or AddaVax (Th1 adjuvant) (**Figure 4C**). We found that pre-exposure to mRNA-LNP did not interfere with the adaptive anti-HA responses induced by Alum- and AddaVax-adjuvanted protein (**Figure 4D and E**). Thus, these data suggest that the inhibition by mRNA-LNPs could be overcome with the use of adjuvants.

### Pre-exposure to mRNA-LNP decreases antigen levels

Since the protein-based adjuvanted vaccine’s efficacy was not affected by pre-exposure to mRNA-LNP, we hypothesized that a possible mechanism of inhibition with the mRNA-LNP platform might lie with mRNA degradation, translation, etc., limiting the production of the antigen coded by the mRNA. The decreased amount of antigen could lead to lower overall adaptive immune responses. To test whether pre-exposure to mRNA-LNP leads to decreased antigen production, animals exposed to PBS, mRNA-LNP coding for luciferase (Luc) or PR8 HA were injected two weeks later with Luc mRNA-LNPs and imaged using IVIS daily for 7 days (Supplemental Figure 2A). We found that pre-exposure to Luc or PR8 HA mRNA-LNP significantly decreased the signals in comparison to PBS or no-exposure controls (Supplemental Figure 2B-D). Thus, these data suggest that pre-exposure to mRNA-LNPs might affect antigen levels by regulating mRNA half-life, translation, etc.

To bring further evidence for lower levels of antigens being produced after pre-exposure to mRNA-LNP, as a second exposure we used DiI-labeled mRNA-LNP coding for eGFP. This reagent allowed us to detect the cells that picked up the mRNA-LNP (DiI^+^) and translated the mRNA into protein (GFP^+^). Two days post inoculation, we found in the skin and skin draining LNs of mice pre-exposed to mRNA-LNP a decrease of DiI/GFP double positive cell percentages (Supplemental Figure 3). Overall, these data are in concordance with decreased luciferase signal presented above, and further support that pre-exposure to mRNA-LNP might interfere with the efficiency of the subsequent shots.

### Pre-exposure to mRNA-LNPs enhances resistance to heterologous influenza infections but decreases resistance to *Candida albicans*

Innate immune cells are sensitive to inflammatory signals and respond with epigenetic modifications that promote or suppress the subsequent innate immune responses [9]. Subsequently, if such effects are induced by certain vaccines, they can impact susceptibility to heterologous infections. To assess whether pre-exposure to mRNA-LNPs affects susceptibility to heterologous infections, we exposed mice to PBS or mRNA-LNP coding for eGFP. Two weeks later, we inoculated the mice with either a sublethal dose of influenza intranasally or *Candida albicans* intravenously. Disease progression was monitored by taking the weight of the mice daily (**Figure 5A**). We found that mice pre-exposed to mRNA-LNP showed significantly higher resistance to influenza infection and lost less weight than the PBS-exposed mice (**Figure 5B**). We also found significantly lower viral loads in the lungs of the mRNA-LNP-pre-exposed mice seven days post inoculation (**Figure 5C**). Flow cytometry analysis of the lungs before and seven days post infection did not reveal major changes in innate immune cell composition (Supplemental Figure 4). In contrast, the mRNA-LNP exposed mice showed significantly diminished resistance towards *Candida albicans* infection. They lost significantly more weight (**Figure 5D**) and we detected approximately a log higher CFU counts in their kidneys (**Figure 5E**), which are the target organs in this mouse model of disseminated candidiasis. Thus, these data suggest that pre-exposure to mRNA-LNPs might alter innate immune fitness.

**Figure 5.**
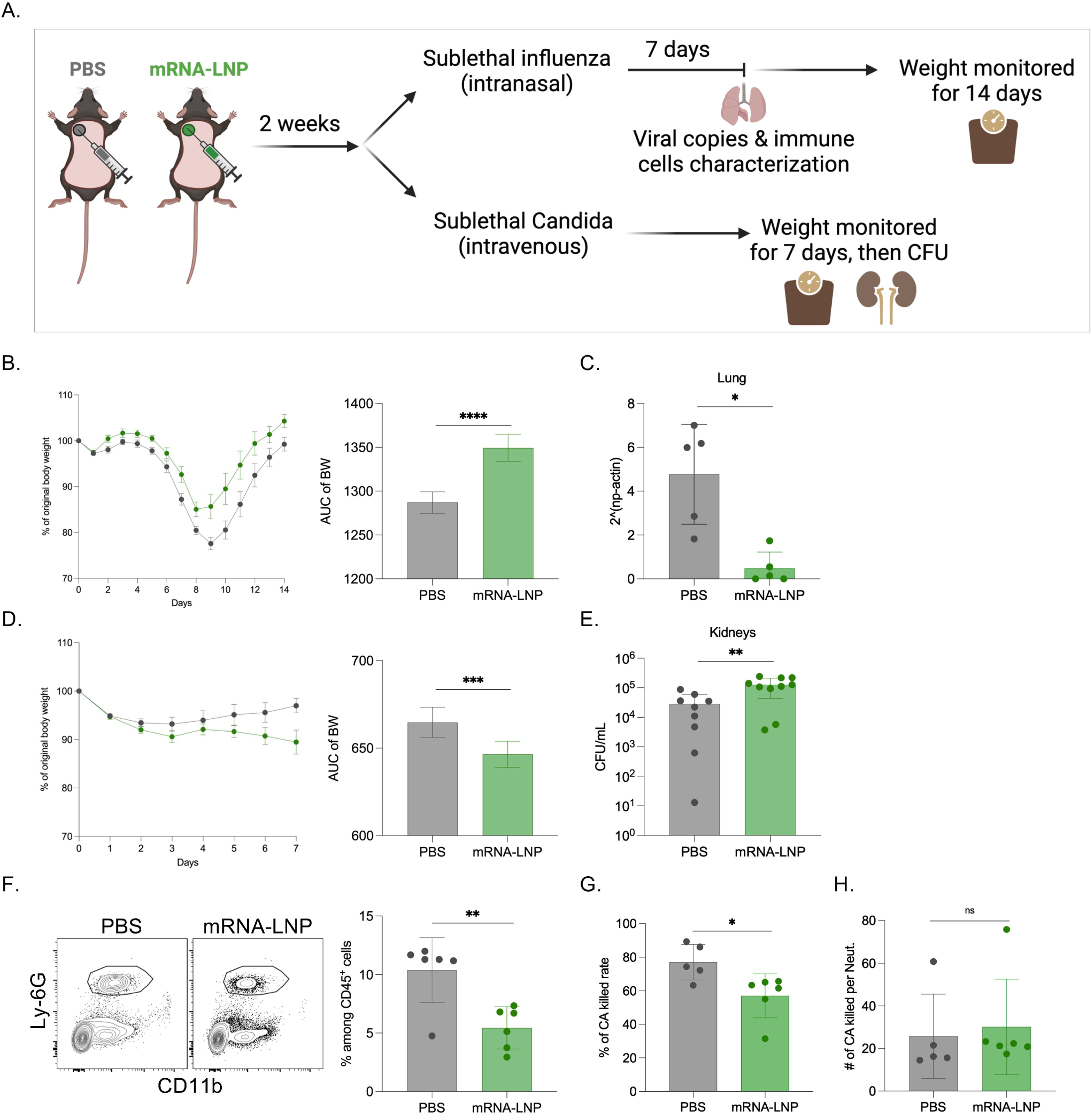
Innate immune fitness is altered with pre-exposure to mRNA-LNP. **A**). Experimental model. Two weeks post-exposure to PBS or eGFP mRNA-LNP the animals were infected with sublethal doses of PR8 HA influenza or *Candida albicans*. The weights and other attributes were monitored as depicted. **B**). Percent of body weight drop after PR8 HA influenza infection, and the corresponding AUC changes. **C**). Viral copies in the lungs. **D**). Percent of body weight drop after *Candida* infection, and the corresponding AUC changes. **E**). *Candida* CFU numbers in the kidneys. **F**). Representative flow plots for neutrophils (gated on live cells/CD45^+^/Ly-6G^+^CD11b^+^) in the PBMCs of PBS or mRNA-LNP exposed mice, and the summary bar graph. **G**). *Candida albicans* (CA) killed rate in a candidacidal assay. **H**). Number of CA killed per neutrophil. Each dot represents a separate mouse. The data were pooled from 2-3 independent experiments. Welch’s t test was used to establish significance. ns = not significant. *p<0.05, **p<0.005, ***p<0.0005, ****p<0.0001.

Next, we set out to determine the possible mechanism by which pre-exposure to mRNA-LNP leads to defective *Candida* clearance. Neutrophils constitute the first line of innate defense and play an important role in fighting fungal infections [16,17]. Therefore, we hypothesized that exposure to mRNA-LNP might alter the neutrophils’ candidacidal properties. To test this, we set up a candidacidal assay [18–20], for which, we used peripheral blood as a source of neutrophils. Flow cytometry analysis of the blood prior the assay revealed unexpectedly a significantly lower neutrophil percentages (**Figure 5F**) in the mRNA-LNP pre-exposed mice, with a concomitant increase in B cells, while no change in DCs and monocytes (Supplemental Figure 5). In line with the decreased neutrophil percentages, we recovered higher *Candida* CFUs from cultures with PBMCs from mRNA-LNP exposed animals (**Figure 5G**). However, the per cell basis normalized candidacidal activity showed no statistical difference between the groups (**Figure 5H**). Thus, these data support that pre-exposure to mRNA-LNP has a profound long-term effect on white blood cell counts and decreases neutrophil percentages but might not interfere with their function. Altogether, the neutropenia reported here might account for lower *in vivo* resistance to *Candida* infection.

### Immune changes induced by pre-exposure to mRNA-LNP can be inherited

Transmission of immune traits to the next generation in vertebrates has been recently reported [21–23]. In humans, lower overall mortality has been reported in infants whose fathers had been vaccinated with BCG [24], and maternal SARS-CoV-2 infection has been associated with increased cytokine functionality and nonspecific immune imprinting in neonates [25]. Finally, trained immunity has been shown to be transmitted in newborn infants of hepatitis B virus-infected mothers [26]. Since the mRNA-LNP platform is highly inflammatory, and we observed that pre-exposure to it enhances resistance to heterologous infection to influenza, we hypothesized that some of these traits might be inherited by the offspring. To test our hypothesis, we immunized adult WT B6 male and female mice intradermally with 10 μg of mRNA-LNP coding for influenza PR8 HA, as we previously described [6]. Two weeks post-immunization, the mice were screened for successful immunization by anti-HA ELISA (Supplemental Figure 6A-B) and then mated (**Figure 6A**) as follows: immunized males with unimmunized (DI; dad immunized) or immunized females (DMI; dad-mom immunized), and immunized females with unimmunized males (MI; mom immunized). Non-immunized males mated to non-immunized females served as controls (DMN; dad-mom non-immunized). Offspring from 1^st^, 2^nd^, and 4^th^ litters at eight-ten weeks of age were intranasally inoculated with a sublethal dose of PR8 influenza and weight monitored for 14 days (**Figure 6A**). We found that mice from the 1^st^ litter from the DI group showed significantly better resistance to influenza infection and lost less weight than the litters derived from naïve parents (**Figure 6B**). Mice from MI or DMI groups showed complete protection from weight drop (**Figure 6B**), which was likely in large mediated by passive immunity provided by the maternal anti-HA antibodies (Supplemental Figure 6C). The second litters derived from the DI group were no longer different from those of non-immunized parents (**Figure 6B**). MI litters showed a significant drop in protection, but still above the litters from DMN parents (**Figure 6B**). Interestingly, litters from DMI still showed complete protection from weight loss (**Figure 6B**). With the 4^th^ litters, the DI and DMN mice remained similar, while the MI and DMI mice were comparable but still significantly protected compared to DMN litters (**Figure 6B**). Thus, these data all together support that the immune changes induced by the mRNA-LNP vaccine in parents can be passed down to the offspring, and both male and female mice play an important role in this transmission.

**Figure 6.**
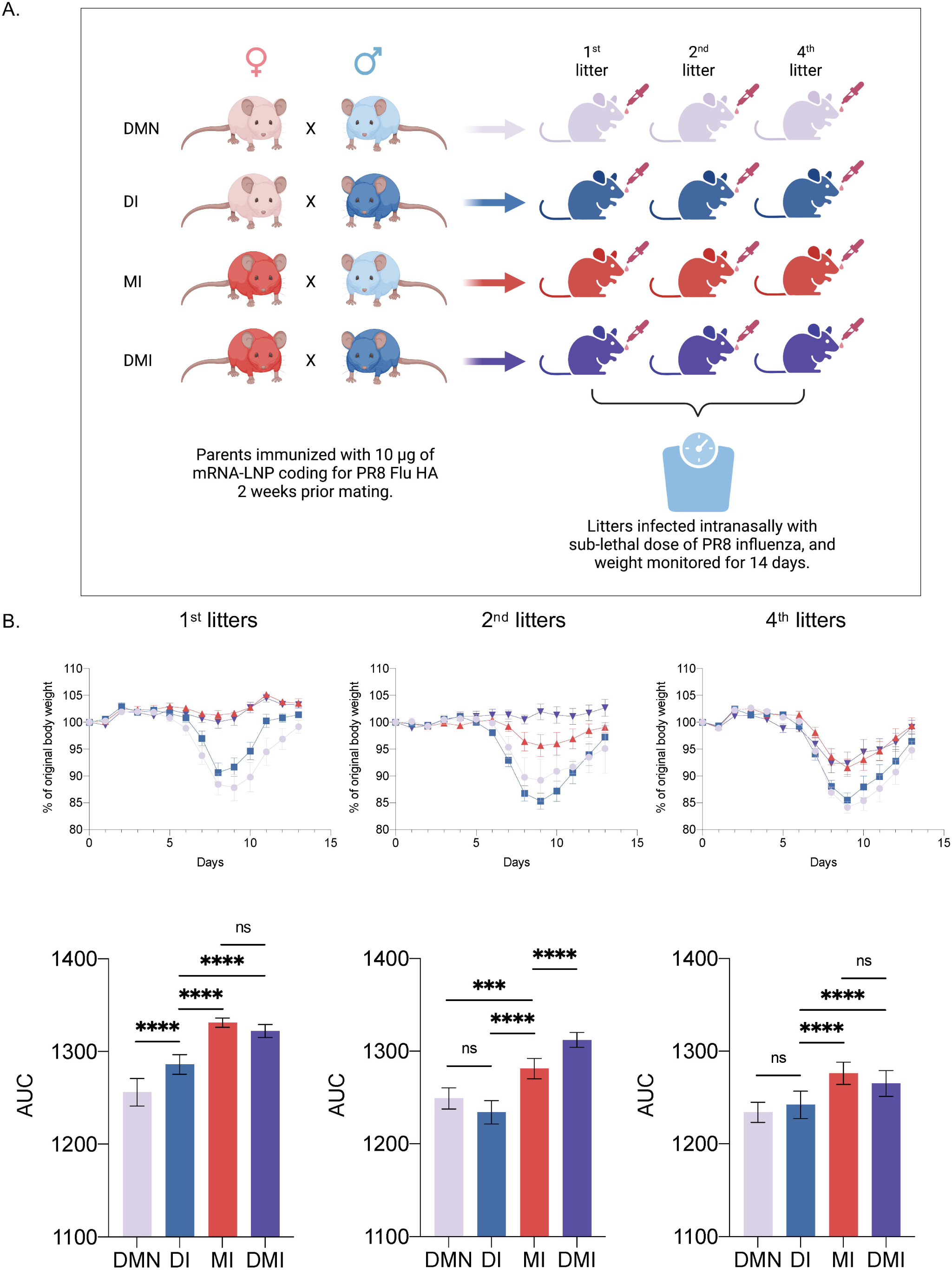
Immune changes induced by the mRNA-LNP platform can be inherited. **A**). Experimental model. Adult WT B6 male and female mice were intradermally immunized with 10 μg of mRNA-LNP coding for influenza PR8 HA. Two weeks post-immunization the mice were mated. Non-immunized males mated to non-immunized females served as controls (DMN; dad-mom non-immunized). Immunized males with unimmunized (DI; dad immunized) or immunized females (DMI; dad-mom immunized), and immunized females with unimmunized males (MI; mom immunized). Offspring from 1^st^, 2^nd^, and 4^th^ litters at eight-ten weeks of age were intranasally inoculated with a sublethal dose of PR8 influenza and weight monitored for 14 days. **B**). Percent of body weight drop after influenza infection (upper), and the corresponding AUC changes (lower). The data are from one experiment using litters from 3 separate mating. The number of mice (female/male) from 1^st^ litters used for influenza challenge was 6/7 (DMN), 8/6 (DI), 3/9 (MI) and 8/7 (DMI); for the 2^nd^ litters 1/3 (DMN), 6/9 (DI), 7/5 (MI) and 6/2 (DMI); for the 4^th^ litters 2/9 (DMN), 8/13 (DI), 9/7 (MI) and 7/8 (DMI). One way ANOVA was used to establish significance. ns = not significant. ***p<0.0005, ****p<0.0001.

## DISCUSSION

The new anti-COVID-19 mRNA vaccines’ immunological effects beyond inducing certain protection against SARS-CoV-2 infection are poorly understood. Based on our earlier studies demonstrating the pro-inflammatory properties of the LNP platform used in these vaccines, we report that pre-exposure to the mRNA-LNP platform has long-term impacts on both innate and adaptive immune responses, with some of these traits being even inherited by the offspring.

The first aim of our study was to assess whether a previous exposure to mRNA-LNPs influences the response to secondary vaccination. Interestingly, we found that indeed the antibody response was inhibited after an earlier administration of mRNA-LNPs. This inhibition of adaptive immune responses was relatively long-lasting, with effects seen for at least 4 weeks, while starting to wane after 8 weeks. Humans receive a 2-dose standard regimen of mRNA-LNP vaccines at 3 to 4 weeks intervals [27] and booster shots at different time points. Our data are strongly supported by recent studies showing that a delay of the second dose of an mRNA vaccine from 3 weeks to 3 months significantly improves the antibody response [28–30]. Indeed, inflammation has been related to a poor responsiveness to vaccination in earlier studies [31], and it is rational to hypothesize that the acute inflammatory side effects of the LNP platform negatively impedes induction of antibody responses during the second dose administration. Increasing the interval between vaccination doses, thus giving time to the immune system to return to homeostasis, is likely to improve the effects of the second dose of the vaccine. Thus, our findings have important implications for improving the schedules of administration for the current mRNA vaccines. However, more studies are needed in the future to assess these effects in more detail.

Whether multiple pre-exposures lead to an even more drastic inhibition of the adaptive immune responses and how much overlap there is between mouse and human data remains to be determined. The inhibition of the adaptive immune responses was more pronounced if the second shot was delivered into the site of pre-exposure. This is in concordance with our recent data showing that the adaptive immune responses with the mRNA-LNP platform are mounted in the lymph node(s), draining the injection site [6], while largely sparing the other secondary lymphoid organs. These data suggest that the inhibition of adaptive immune responses by the mRNA-LNP could be partially mitigated if the follow-up shots are delivered far away from the first injection sites.

A possible mechanism of inhibition might lie with mRNA degradation, translation, etc., limiting the production of the antigen coded by the mRNA. The decreased amount of antigen could lead to lower overall adaptive immune responses. Our preliminary data support this hypothesis. We found using two independent systems that antigen levels are likely lower with the mRNA-LNP platform in mice pre-exposed to mRNA-LNP (Supplemental Figure 2 and 3). If further studies confirm that pre-exposure to mRNA-LNP indeed inhibit subsequent adaptive immune responses through regulation of mRNA degradation and translation, that could indicate that the nucleoside modification step of the mRNA might be less important. The nucleoside modification and removal of double stranded RNAs were aimed to lower innate immune recognition, activation, and IFNα secretion to decrease side effects and prolong the mRNAs’ half-life [1–3]. However, our preliminary data support that the LNP component, likely through its highly inflammatory nature, might at least partially counteract the benefit of nucleoside modification and RNA purification. We recently reported that in the mRNA-LNP-exposed skin, several inflammatory pathways, including the Toll-like-, RIG-1-like- and NOD-1-like receptor signaling pathways were highly upregulated [4]. Thus, LNP and mRNA-LNP can likely directly or indirectly engage receptors to induce interferons [4,7] and reduce subsequent mRNA translation. Pre-exposure to TLR4 ligand significantly decreased RNA translation from mRNA-LNP [32,33]. Altogether these data suggest that the mRNA-LNP platform efficiency might be influenced by pre-exposure to vaccines engaging the TLR/interferon pathways or by pre-existing inflammatory environments such as autoimmune disease or ongoing infections.

Interestingly, using a steady-state antigen-targeting model, we found that pre-exposure to mRNA-LNP also led to inhibition of CD4 T cell responses. We observed a decreased CD4 T cell precursor number in mice exposed to mRNA-LNP, which correlated with reduced expansion. Out-of-sequence (prior TCR engagement) stimulation with type I interferon and other cytokines of naïve T cells can lead to their inhibition and apoptosis [34,35]. Whether exposure to mRNA-LNP restricts CD4 T cell responses using a similar mechanism, remains to be explored.

Our data support that the inhibition of the adaptive immune responses could be overcome with the use of adjuvants, including Th1 (AddaVax) and Th2 (Alum) adjuvants. This is a highly relevant finding from a human health perspective. The lack of interference with protein/subunit vaccines is reassuring that their efficacy will not be hindered by the pre-exposure to the mRNA-LNP platform. Indeed, it has been also shown that combination of mRNA vaccines with adenovirus-based anti-SARS-CoV-2 vaccines may actually even improve the serological responses and protection [36–38]. However, whether the effectiveness of live attenuated viral vaccines, such as influenza is affected, remains to be determined.

We found that mice pre-exposed to mRNA-LNP showed enhanced resistance to influenza, which correlated with lower viral loads in the lungs. Why mice are better protected from influenza remains to be determined. We observed no significant changes in the innate immune cell composition of the lungs from PBS and mRNA-LNP pretreated mice before and 7 days post-infection. One potential mechanism by which the exposure to mRNA-LNP could confer enhanced protection from influenza might lie with RNA biology regulated by inflammatory cytokines such as interferons [34,39]. As we showed, antigen levels were lower with the second shot. Thus, the virus replication may be inhibited through a similar mechanism, especially since influenza is an RNA virus. In contrast to influenza, mice exposed to mRNA-LNP showed defective clearance of disseminating *Candida albicans* infections. The failure to efficiently clear *Candida* correlated with neutropenia in the blood. What led to changes in neutrophil counts after exposure to mRNA-LNP remains to be determined. Interferons highly induced by this platform [7] might interfere with protection [35] and are also known to inhibit hematopoiesis [40–42]. Altered hematopoiesis might have contributed to neutropenia. Severe aplastic anemia (affecting the hematopoietic stem cells and characterized by neutropenia and thrombocytopenia)-, cytopenia- and thrombocytopenia cases have all been reported with this platform in humans [43–49]. One small human study (56 volunteers) that monitored the immunological changes induced by this vaccine up to 21 days of the second shot reported a significant fluctuation in white blood cell numbers after the second injection of mRNA-LNP [11]. The white blood cell numbers seemed to normalize a week after the injection. What led to these sudden changes in white blood cell counts and whether further and more stable changes can be observed months after vaccination remains to be determined. It also remains to be defined how much overlap exists between our data and the still-to-be peer-reviewed human observation on innate and adaptive reprogramming with this platform [10]. If our data can be translated to humans, it is anticipated that people might present with an altered incidence of certain infections. In line with this, a recent retrospective study found that vaccinated people might show a higher risk of infection than unvaccinated individuals 9 months post-vaccination [50]. A potential sign of an immunosuppressed state comes from reports of viral reactivations [51–55] and suspected infections in open-heart surgeries that could not be controlled even with long-term antibiotic treatments, resulting in several deaths [56]. A surge in candidiasis, aspergillosis, and mucormycosis cases associated with COVID-19 has been reported [57–59]. Whether any of these cases could be attributed to exposure to the mRNA-LNP vaccines remains to be determined. Further research will also be needed to establish the long-term effect of multiple mRNA-LNP shots on immune fitness of humans and mice.

In humans, many vaccines, including the mRNA-LNP vaccines are administered intramuscularly for ease of use [60]. The preclinical animal intradermal and intramuscular vaccine studies revealed induction of similar adaptive immune responses, with slightly better responses with the intradermal delivery [61,62]. The inflammatory reactions in nature and magnitude induced by the mRNA-LNP platform are independent of the delivery route [4]. They are characterized by robust and transient neutrophil influx and activation of multiple different inflammatory pathways and the production of inflammatory cytokines such as IL-1β and IL-6. Thus, it is unlikely that the immune reprogramming observed here would be limited to intradermal exposure, especially since data from mRNA-LNP-vaccinated individuals collaborate our findings [10]. The vaccine doses used in mouse and human studies differ significantly, and human vaccines doses adjusted to weight are much lower than in mice [63]. In our studies, we used doses at the lower end of the amounts used for mouse vaccine studies [61]. While we did not perform a wide dose range for our studies, we observed that pre-exposure to 2.5 μg of empty LNPs and 2.5 μg mRNA-LNPs (2.5 μg refers to the amount of mRNA, which is complexed with LNPs at ∼1:20 or 30 weight ratio, i.e., ∼50 or 75 μg of LNP) resulted in similar inhibition of adaptive immune responses. Thus, while dose might play a role in the magnitude and length of inhibition, our data combined with human data support that the LNPs/mRNA-LNps have similar effects in a wide dose range.

A very interesting observation was that the improved heterologous protection induced by mRNA-LNPs on influenza infection was successfully passed down to the offspring. A number of recent studies reported evidence for transmission of either trained immunity or tolerance across generations in mice [23,64], although not all of them [65]. Our independent study initiated and performed before these data became public seem to support the existence of transgenerational inheritance of immune traits. The highly inflammatory properties of the mRNA-LNP platform might have induced the inherited changes, and it would be very important to determine whether if any such immune inheritance may be observed in humans vaccinated with mRNA vaccines. Cross-generational protection of infants of parents vaccinated with BCG has recently suggested in epidemiological studies [24]. Whether these changes in innate immune genes could also directly or indirectly affect adaptive immune responses remains to be determined. Our experimental platform was able to detect the innate contribution of males. The passively transferred maternal antibodies largely masked the innate female contribution. Nevertheless, we observed that the 2^nd^ litters from the DMI group outperformed the MI ones. These data suggest that innate immune traits inherited from the father could provide essential advantages in protection even in the presence of maternal-derived antibodies. However, since the 2^nd^ litters from the DI group were no better than the litters from DMN, but the DMI litters outperformed the MI litters, these data suggest that an immune female counterpart might further boost the benefits provided by the immune traits inherited from males. Nevertheless, the overall protection levels fell across the board with later litters, suggesting that such heterologous effects do not persist for the entire life of an animal. While here we focused our attention on whether the offspring are more resistant to influenza, it will be important to define how long after immunization the parents can still pass down the acquired immune traits and whether the offspring’s resistance to fungal infections decreases, and whether the innate immune changes alter the adaptive immune responses. The mechanism of inheritance also remains to be determined. Likely, it is partially mediated through DNA methylation changes that interferons and other inflammatory cytokines might have induced in this case. DNA methylation-based mode of inheritance has recently been proposed with the transgenerational inheritance observed with pre-exposure to different pathogens [64].

In conclusion, we describe important immunological properties of the mRNA-LNP platform used in mRNA vaccines against COVID-19. These findings have important biological and clinical implications. First, our data show that the administration of mRNA-LNPs inhibits humoral responses to a second dose of the vaccine for at least several weeks: this finding is highly relevant from a human health perspective as it suggests that a second dose of a mRNA vaccine may be more effective if given at a later timepoint and different location than currently used. This conclusion is strongly supported by recent human studies suggesting that indeed the delay of the second dose of the Pfizer/BioNTech vaccine leads to a better humoral and cellular response [28,29]. Second, our study suggests altered heterologous protection against fungal and viral infections by the mRNA-LNPs platform. Third, our study also shows the capacity of these vaccines to transmit protection trans-generationally, thus supporting the concept of Lamarckian inheritance in mammals [66]. However, our study only partially opens the door towards understanding the various immunological effects of mRNA-LNP platform. Considering the broad exposure of a large proportion of human populations to vaccines based on this novel technology, more studies are warranted to fully understand its overall immunological and physiological effects. Determining this platform’s short and long-term impact on human health would help optimize it to decrease its potentially harmful effects.

## ACKNOWLEDGEMENTS

We would like to thank Dr. Scott Hensley (UPenn) for the PR8 HA influenza stock, and Drs. Barney Graham and Masaru Kanekiyo at NIH for the recombinant HA protein. B.Z.I. is supported by NIH R01AI146420, R01AI146101 and departmental start-up funds. We thank Drs. Tim Manser at TJU and Mihai Netea at Radboud University for critically reading and editing the manuscript. Research reported in this publication utilized the Flow Cytometry and Laboratory Animal facilities at the Sidney Kimmel Cancer Center at Jefferson Health and was supported by the National Cancer Institute of the National Institutes of Health under award number P30CA056036. Figures were generated using BioRender.

## AUTHOR CONTRIBUTION

Z.Q. performed experiments and analyzed the data. A.B. and C.H. helped with the revision. B.Z.I. conceptualized the study, interpreted the data, and wrote the manuscript.

## DECLARATION OF INTERESTS

Authors declare no conflict of any sort.

---

**Supplemental Figure 1. Gating strategy for HA-specific GC B cells**

**Supplemental Figure 2. Pre-exposure to mRNA-LNPs decreases antigen levels. A**). Experimental model. Balb/c mice were pre-exposed to PBS, Luc mRNA-LNPs or PR8 HA mRNA-LNPs and imaged using IVIS 6 hours (0.25 day) post inoculation and then every day for 7 days. Two weeks later all the animals were injected in the same spot with Luc mRNA-LNPs and the luciferase signal monitored similarly to the first exposure. **B**). Relative total flux with time. **C**). Data from **B** presented as AUC. **D**). Total flux values (background subtracted) of each mouse at different time points are shown as log_10_. X marks mice where the signal was below detection. Data from two separate experiments pooled. One way ANOVA was used to establish significance. ns = not significant. ***p<0.0005, ****p<0.0001.

**Supplemental Figure 3. Pre-exposure to mRNA-LNPs leads to overall decrease of antigen levels. A**). Experimental model. Animals were shaved and intradermally inoculated in the left upper spot with either PBS or mRNA-LNP coding for HA. Two weeks later the same areas were injected with mRNA-LNP-DiI coding for eGFP. The injected skin (2cm^2^) and skin draining lymph nodes were harvested 2 days later and the eGFP and DiI signals determined using flow cytometer. **B**). Representative flow plots and summary graph on eGFP^+^DiI^+^ population of skin DCs (MHCII^+^ CD11c^+^) after gating on live cell/Ly-6G^-^/CD64^-^. Naïve mice were not injected with mRNA-LNP-DiI coding for eGFP. **C**). Representative flow plots and summary graphs on eGFP^+^DiI^+^ population of SDLNs CD45^-^ cells, macrophages (Mac., CD64^+^), B cells (MHCII^+^CD11c^-^) and mDCs (MHCII^high^ CD11c^mid^). Each dot represents a separate mouse. The data are from one experiment and are shown as mean ±SD. Welch’s t test was used to establish significance. ns = not significant. *p<0.05.

**Supplemental Figure 4. Characterization of lung immune cells before and after influenza challenge in PBS or mRNA-LNP pre-exposed mice**. Summary graphs of levels of lung immune cells from pre- and post-influenza infected mice which were pre-exposed to PBS or mRNA-LNP for 2 weeks. Each dot represents a separate mouse. The data were pooled from two experiments. Welch’s t test was used to establish significance. ns = not significant. *p<0.05.

**Supplemental Figure 5. PBMC levels after pre-exposure to PBS or mRNA-LNP**. Summary graph of total cell number and major categories of cells among CD45^+^ cells in PBMCs from mice pre-exposed to either PBS or mRNA-LNP for 2 weeks. Each dot represents a separate mouse. The data were pooled from two experiments. Welch’s t test was used to establish significance. ns = not significant. *p<0.05.

**Supplemental Figure 6. Anti-HA antibody levels in parents and litters. A**). Anti-HA antibody levels in parents 2 weeks post inoculation. **B**). Anti-HA antibody levels in parents 2 weeks and 28 weeks post inoculation. **C**). Anti-HA antibody levels in the 1^st^ litters prior- and 4 weeks post-infection.

## MATERIALS AND METHODS

### Ethics statement

Institutional Care and Use Committee at Thomas Jefferson University approved all mouse protocols. Protocol number: 02315.

### Mice

WT C57BL/6J and Balb/c mice were purchased from Jax and bred in house. Balb/c mice were only used for the IVIS experiments. All experiments were performed with 8– 12 weeks old female and male mice. Mice were housed in microisolator cages and fed autoclaved food.

### mRNA-LNPs

For our studies, we used an LNP formulation proprietary to Acuitas Therapeutics described in US patent US10,221,127. These LNPs were previously carefully characterized and widely tested in preclinical vaccine studies in combination with nucleoside-modified mRNAs [62,67]. The following, previously published mRNA-LNP formulations were used: PR8 HA mRNA-LNP, Luc mRNA-LNP, eGFP mRNA-LNP-DiI and empty LNPs.

### Infectious agents

A/Puerto Rico/8/1934 influenza stock was a generous gift from Dr. Scott Hensley (University of Pennsylvania). *C. albicans* strains used in this study were previously described [68]. The work with the infectious agents was performed in a BSL2 laboratory and approved by the Institutional Biosafety Committee.

### Intradermal immunization

Intradermal immunizations were performed as we previously described [4,6]. Briefly, the hair of the site of injections was wet shaved using Personna razor blades. The mice were then injected intradermally (upper left area on the back) with 2.5 μg/spot mRNA-LNPs or empty LNPs in 20 μl PBS or equivalent volume of PBS as 1^st^ shot. The 2^nd^ shot of 2.5 μg/spot mRNA-LNPs was administered 2, 4 or 8 weeks after the 1^st^ shot at either the same or opposite site on the upper back area. To test for platform specificity, 5 μg HA protein combined with either Alum (InvivoGen) or AddaVax (InvivoGen) at 1:1 (v/v) ratio was intradermally injected at the same site two weeks after the 1^st^ mRNA-LNP shot. For the inherited immunity experiment the parents were injected with 10 μg (2.5 μg/spot; 4 spots) of mRNA-LNP coding for PR8 HA or with the corresponding volume of PBS [4,6].

### Steady-state antigen targeting

These experiments were performed as previously described [14,15] with slight modifications. Briefly, mice received intravenous (i.v.) transfer of CFSE-labeled, congenically marked 5×10^5^ TEα cells 1 day before or 13 days after the intradermal exposure to mRNA(eGFP)-LNP or PBS. One μg of anti-Langerin-Eα or PBS were administered i.v. on day 14. Mice were then sacrificed 4 days later and the skin draining lymph nodes (axillary and brachial) harvested. Single-cell suspensions were stained with fixable viability dye (Thermo Fisher), CD4 (GK1.5), and CD90.1 (OX-7). The antibodies were purchased from BioLegend.

### Recombinant hemagglutinin (HA) protein labeling

Recombinant HA protein (rHA) was a kind gift from Drs. Barney Graham and Masaru Kanekiyo at NIH. Molecular Probes Alexa Fluor 647 Protein Labeling kit (A20173, Invitrogen) was used to label rHA following the product user guide.

### Characterization of B cell responses

At day 14 post-injections, the mice were sacrificed and the skin draining lymph nodes (axillary and brachial) harvested. Single-cell suspensions were generated using mechanical disruption through cell strainers. The cells were stained with B cell panel consisting of dump (fixable viability dye, F4/80, CD11b), CD38 (90), B220 (RA3-6B2), CD138 (281–2), GL-7 (GL-7), Sca-1 (D7), IgD (11-26c.2a), IgM (RMM-1) and AF647-labeled rHA. A gating strategy previously published (Supplemental Figure 1) was used to define GC B cell populations [6]. The GC B cell percentages were normalized between experiments and are shown as relative values. The normalization was performed as follows. The mean GC % of all the samples from one experiment was used to divide each sample GC % value (relative value). The antibodies were purchased from BD Biosciences, Biolegend or Tonbo Biosciences. The stained samples were run on Fortessa (BD Biosciences) and the resulting data analyzed with FlowJo 10.

### *In vivo* challenge with PR8 influenza or *Candida albicans*

The doses of influenza or *C.albicans* used in these studies were previously described [64,69,70]. For viral infection, mice were anesthetized by intraperitoneal injection with a mixture of Xylazine/Ketamine and inoculated intranasally with 200 TCDI_50_ PR8 influenza virus. For fungal challenge, mice were intravenously injected with 3∼4×10^4^ CFU *C.albicans*. Subsequently the mice were monitored daily for distress and weight loss. The weight loss data are presented as percent of original body weight.

### Characterization of immune cells in the lungs

Mice were injected intradermally with 2.5 μg mRNA(GFP)-LNP or PBS at the upper left site of the back. Two weeks later, the mice were intranasally inoculated with 200 TCID_50_ PR8 influenza virus. Seven days post infection, mouse lungs were collected the mice were sacrificed and the lungs harvested. Single-cell suspension was prepared as we previously described [71]. Briefly, lungs were minced, digested with Collagenase XI (Sigma-Aldrich) and DNase (Sigma-Aldrich), and passed through cell strainer. The red blood cells were lysed with ACK lysis buffer (Fisher Scientific). The cells were stained with fixable viability dye (Thermo Fisher), CD45 (30 F11), Ly-6G (1A8), CD11b (M1/70), MHCII (M5/114.15.2), CD24 (M1/69), Ly-6C (HK1.4), CD64 (X54-5/7.1), and CD11c (N418).

### Quantification of viral burden

Mice were injected intradermally with 2.5 μg mRNA(GFP)-LNP or PBS at the upper left site of the back. Two weeks later, the mice were intranasally inoculated with 200 TCID_50_ PR8 influenza virus. Seven days post infection, mouse lungs were collected, and flash frozen prior to storage at -80 °C. RNA was prepared using the E.Z.N.A. HP Total RNA Kit (Omega Bio-tek) following the manufacturer’s instructions. One μg of RNA was reverse-transcribed using iScript Reverse Transcription Supermix for RT-PCR (Bio Rad), following the manufacturer’s instructions. Quantitative PCR was performed with iTaq Universal SYBR Green Supermix (Bio Rad), following the manufacturer’s instructions. Relative viral load was measured by ΔCT of PR8 influenza virus nucleoprotein (NP) using mouse β-Actin as a reference gene. Forward (5’) Pr8 NP: CAGCCTAATCAGACCAAATG, Reverse (3’): TACCTGCTTCTCAGTTCAAG. Forward (5’) β-Actin AGATTACTGCTCTGGCTCCTAGC and Reverse (3’): ACTCATCGTACTCCTGCTTGCT [72,73].

### Quantification of fungal burden

On day 7 post *C.albicans* challenge, mice were sacrificed and the kidneys harvested. The organs were then homogenized in PBS, and the homogenates were diluted and used to inoculate YPAD agar plates. After overnight incubation at 30°C, the colonies were manually counted, and the CFU/ml organ calculated.

### Candidacidal assay

Mice were injected intradermally with 2.5 μg mRNA(eGFP)-LNP or PBS at upper left side of the back. Two weeks later, mice were sacrificed, and the blood samples collected for PBMC isolation and serum collection. To isolate PBMCs, the blood was first mixed with 0.5 M EDTA at 9:1 ratio and then with equal amount of PBS. The mixtures were then centrifuged at 4C° at 300 g for 6 minutes. The pellets were resuspended in ACK and incubated for 5 minutes, then washed with staining media twice. The resulting pellets were resuspended in 300 μl staining media and counted. 4×10^5^ cells/well were seeded in a 96-well flat bottom plate and incubated at 37 C° with 5 % CO_2_ for 1h. *C.albicans* (CA) were opsonized with mouse serum (10% final concentration), 100 μl of which containing 1.6×10^3^ CA was added into each well with/out PBMCs. After 2h incubation, the supernatant was collected and the wells were washed with ddH_2_O, which were combined and used for streaking YPAD agar plate [18–20]. The CFU was read after overnight incubation at 30 C°.

### Serum collection

To collect the serum, the blood samples were left to coagulate at room temperature for 30 minutes, and then centrifuged at 4 C° at 13,500 g for 18 minutes.

### Characterization of white blood cells

PBMCs were stained with the following reagents and antibodies: fixable viability dye (Thermo Fisher), CD45 (30 F11), CD11b (M1/70), Ly-6G (1A8), CD64 (X54-5/7.1), CD11c (N418) and MHCII (M5/114.15.2). The antibodies were from BioLegend, BD Biosciences and Tonbo Biosciences.

## ELISA

Nunc Immuno 96 well plates (Fisher Scientific) were coated with 1μg/ml (50 μl/well) HA protein (Sino Biological) diluted in carbonate/bicarbonate buffer (Fisher Scientific) overnight at 4°C or 1 hour at 37 °C. After washing and blocking with TBS for 1 hour the serum samples were diluted and added to the plate. Serially diluted HA-specific monoclonal antibody (Sino Biological) served as standard. Anti-mouse IgG-HRP (1:20,000; Fisher Scientific) in combination with TMB (Fisher Scientific) solution was used for detection. The signals were read at 450 nm using accuSkan FC microplate photometer (Fisher Scientific).

### Analysis of antigen levels

Mice were injected intradermally with 2.5ug mRNA(HA)-LNP or PBS at the upper left site of the back. Two weeks later, the same spot was injected with 2.5ug mRNA(eGFP)-LNP-DiI. Two days post injection, the mice were sacrificed and the skin and skin draining lymph nodes (axillary and brachial) were harvested. Single-cell suspensions were stained with fixable viability dye (Thermo Fisher), anti-CD45 (30 F11), CD11b (M1/70), Ly-6G (1A8), CD64 (X54-5/7.1), CD11c (N418) and MHCII (M5/114.15.2), as we previously described [71]. The antibodies were purchased from BioLegend, BD Biosciences and Tonbo Biosciences.

### *In vivo* bioluminescence imaging

The imaging was performed with IVIS Lumina *XR* system (Caliper Life Sciences). To limit possible interference of melanin in B6 mouse skin with the signal generated by the luciferase activity, we used WT Balb/c mice for these experiments. Mice injected with PBS, mRNA-LNP coding for luciferase or PR8 HA, were intraperitoneally injected with D-Luciferin (Potassium Salt, Goldbio) at the dose of 150 mg/kg. Five minutes later, the mice were anesthetized in a chamber filled with 3 % isoflurane for 1 minute, then transferred to the imaging platform with maintained 2 % isoflurane via gas ports. With the Living Image Software provided by Caliper, the signal was acquired by measuring total flux (photons/sec) for 5 seconds exposure time. The total flux is the radiance (photons/sec/cm^2^/steradian) in each pixel summed or integrated over the region of interest (ROI) area (cm^2^) x 4π. The total flux values were normalized to ensure accurate quantitation and comparability. Briefly, the normalized value (Vn) in each independent experiment was calculated as following:

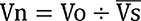

Vo was original total flux value of a given sample at a given timepoint; ^-^V^--^s was the mean of all values from all groups.

### Statistical analyses

All data were analyzed with GraphPad Prism version 9.0.0. Statistical methods used to determine significance are listed under each figure.

## Notes

### Competing Interest Statement

The authors have declared no competing interest.

### Summary of Updates

The revision includes revised Candida data and several extra figures.

