## Supplementary figures and images for "Pre-exposure to mRNA-LNP inhibits adaptive immune responses and alters innate immune fitness in an inheritable fashion"

### Suppl. Figures

Suppl. Figure 1

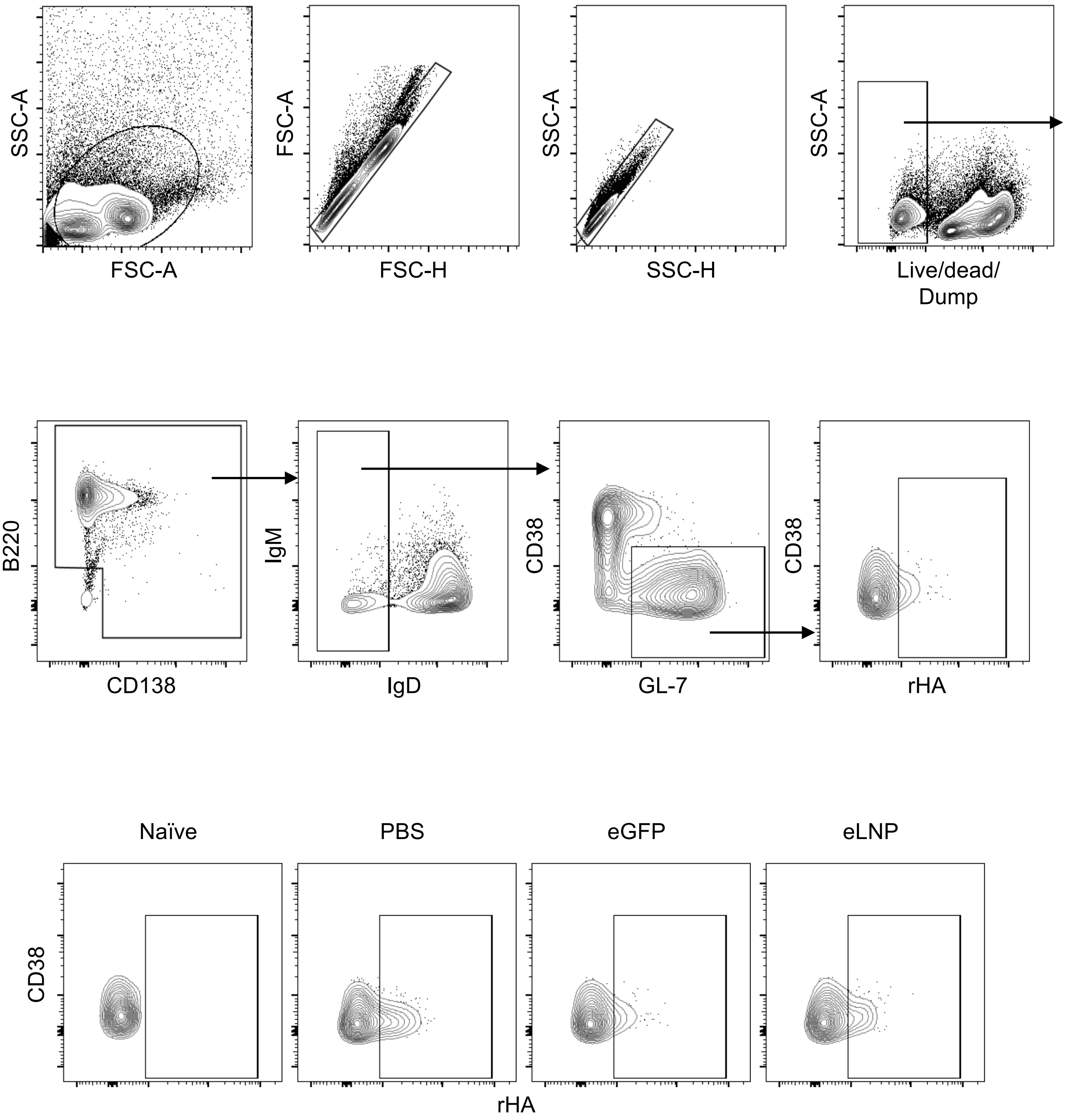

Suppl. Figure 2

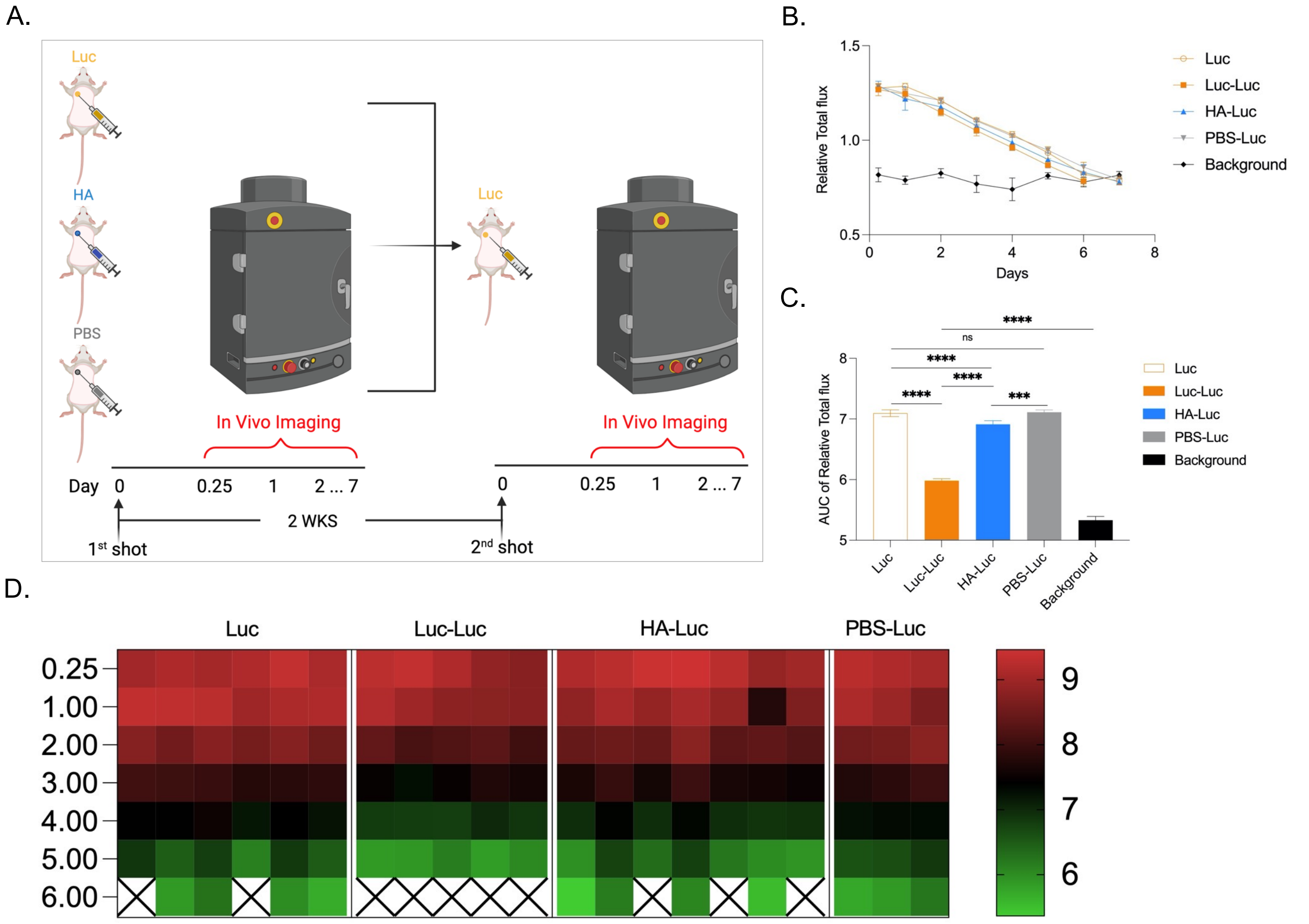

Suppl. Figure 3

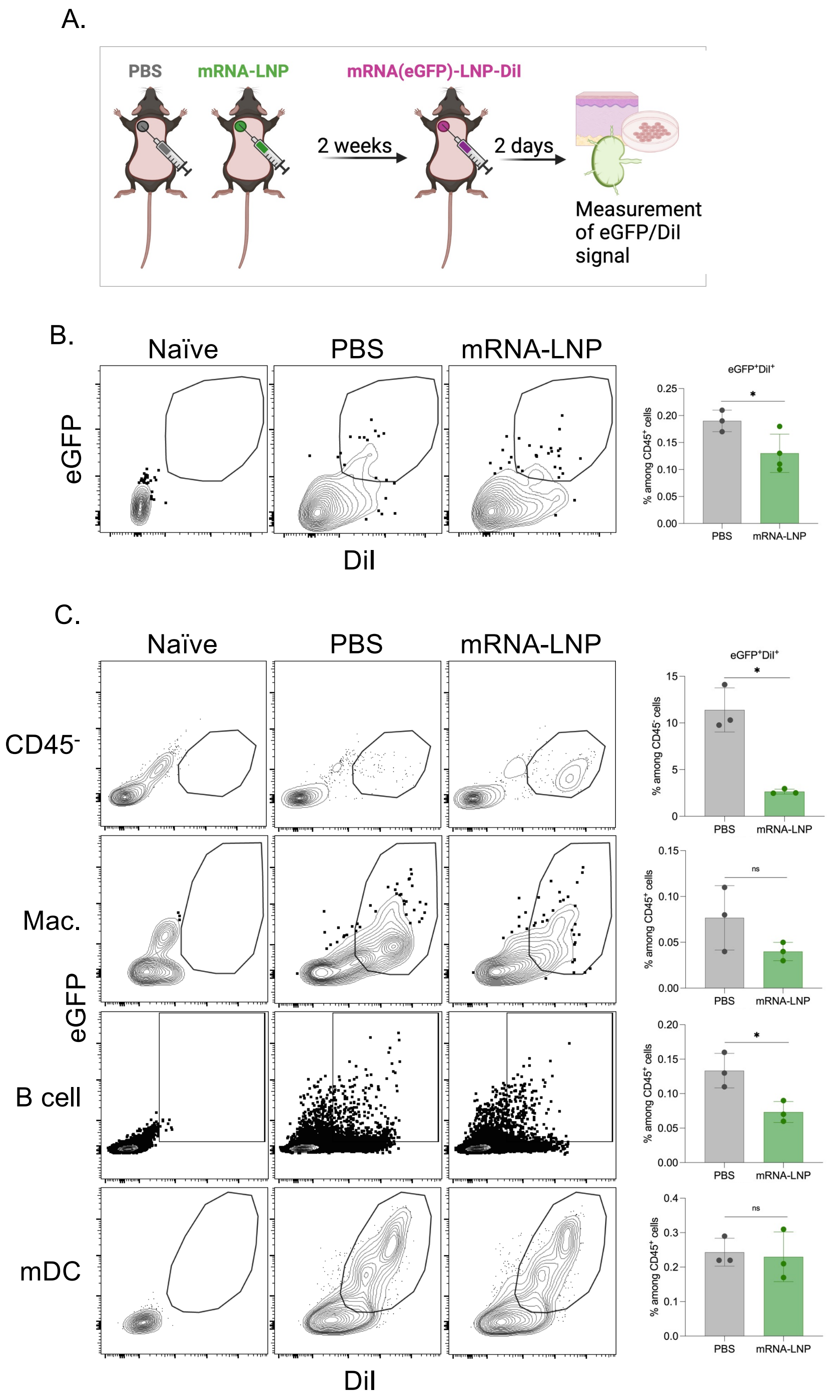

Suppl. Figure 4

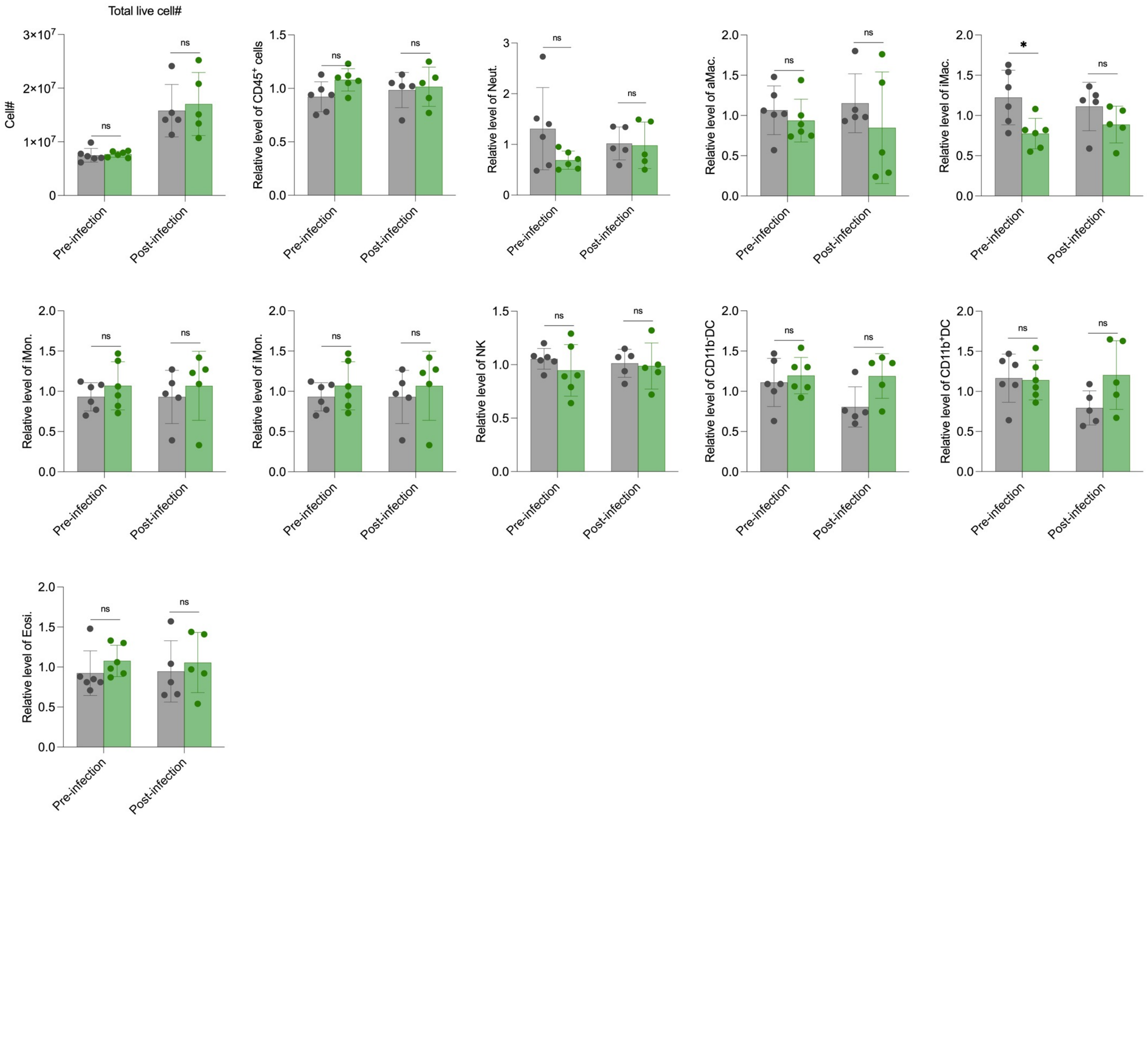

Suppl. Figure 5

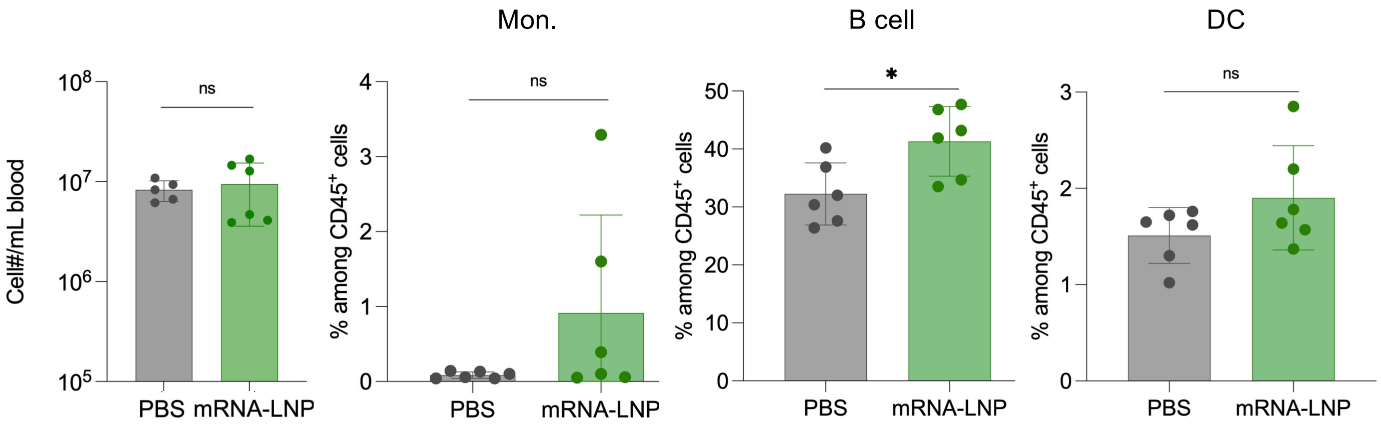

Suppl. Figure 6

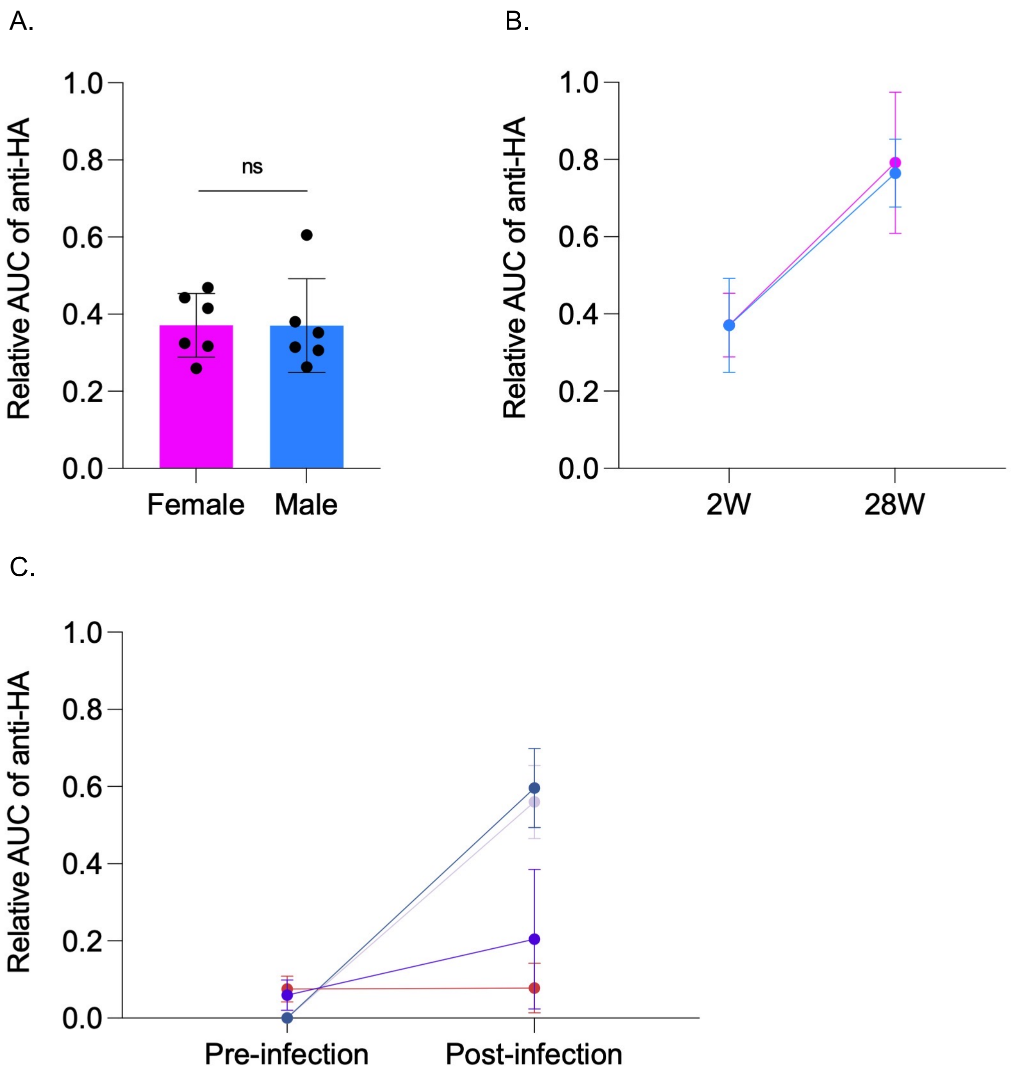
